# Long read sequencing and assembly of wild diploid relatives and cultivars in support of banana breeding programs

**DOI:** 10.1101/2024.09.03.610537

**Authors:** Andrew Chen, Xuefeng Zhou, Keith Decker, Ericka Havecker, Zijin Du, Allan F. Brown, Elizabeth A. B. Aitken, Shah Trushar, Brigitte Uwimana, Chelly Hresko, Rony Swennen, Rebecca F. Lowdon

**Affiliations:** School of Agriculture and Food Sustainability, The University of Queensland, Brisbane, QLD. 4067, Australia. A.C.; E.A.B.A.; Bayer Crop Science, Chesterfield, MO, USA. X.Z., K.D.; E.H.; Z.D.; C.H.; International Institute of Tropical Agriculture, Arusha P.O. Box 447, Tanzania.; International Institute of Tropical Agriculture, Nairobi P.O. Box 30709-00100, Kenya. S.T.; International Institute of Tropical Agriculture, Kampala P.O. Box 7878, Uganda. B.U.; Division of Crop Biotechnics, Laboratory of Tropical Crop Improvement, Katholieke Universiteit Leuven, 3001 Leuven, Belgium

**Keywords:** Banana breeding, PacBio HiFi, chromosome-scale genome assembly, banana, *Musa acuminata*, *Fusarium oxysporum* f. sp. *cubense*, Tropical race 4, Fusarium wilt of Banana

## Abstract

Banana is an important fruit and staple crop, which is vital for food and income security in developing countries. A genomics initiative was undertaken as part of the Global Alliance against TR4 (Tropical Race 4 of Fusarium Wilt of Banana) Alliance, with Bayer Crop Science, International Institute of Tropical Agriculture, and University of Queensland to collectively accelerate the breeding efforts in Eastern Africa. The goal of this project is to develop genomics resources to improve banana breeding, productivity, and quality traits amid a changing climate and shifting disease pressures for breeding. Here, we present the PacBio HiFi sequencing, assembly, and comparative analysis of seven genomes of wild and edible diploid banana accessions belonging to subspecies considered important progenitors to cultivated banana varieties and part of breeding programs. This banana genome resource will help power genome editing, a pangenome effort, and conventional breeding programs for germplasm improvement.

## Introduction

Bananas (*Musa* spp.) are an important horticultural crop that is served as either staple food or fruit, providing sustenance in the form of essential nutrients to millions of people around the world. In 2021, 12 million hectares were under banana cultivation, producing 170 million tonnes of bananas (FAO, 2023). Approximately 26.4 million tons of bananas, 15.5% of total production, are exported each year making it the most exported fruit in the world (FAO, 2023). The export market is dominated by Cavendish varieties. Its primary producers are based in Latin America and the Caribbean, which account for 80% of the export market estimated at 11 billion USD (FAO, 2023). 84.5% of bananas are consumed locally and are grown by small holder farmers (Solidaridad, 2023).

Cultivated bananas including plantains, highland cooking bananas, and Cavendish bananas are mostly derived from hybridizations between *Musa acuminata* (A genome, 2n = 22) and *Musa balbisiana* (B genome, 2n = 22) (Rouard et al., 2018). Increasingly, genome assemblies shed light on the genome structure and functional divergence of diploid and polyploid *Musa* spp. (D’Hont et al., 2012; Rouard et al., 2018; Wang et al., 2019; Martin et al., 2020b; Belser et al., 2021; Huang et al., 2023; Liu et al., 2023; Li et al., 2024). However, breeding programs require sequence knowledge of specific breeding accessions to facilitate marker-assisted breeding and genomic selection. The public availability of genome assemblies of parental accessions important to breeding programs are currently lacking.

Like other clonally propagated tropical crops, bananas are bred using a few elite cultivars and thus are subject to high disease and pest pressures in the field (Jones, 2018), with significant outbreaks of diseases attributed to bacterial (*Xanthomonas*) wilt (Tushemereirwe et al., 2004; Biruma et al., 2007; Reeder et al., 2007; Smith et al., 2008; Mbaka et al., 2009; Tripathi et al., 2009; Carter et al., 2010), fungal pathogens such as *Pseudocercospora fijiensis* (Churchill, 2011; Arango Isaza et al., 2016; Alakonya et al., 2018) and *Fusarium oxysporum* f. sp. *cubense* (Rutherford, 2001; Ploetz, 2015; Dita et al., 2018) as the respective causal agents for black sigatoka and Fusarium wilt, other crippling disease infections caused by the banana bunchy top virus (Dale, 1987; Su et al., 2003; Niyongere et al., 2012; Kumar et al., 2015), root-knot nematode (Speijer et al., 1999; Gaidashova et al., 2009; Onkendi et al., 2014), and banana weevil borer (Rukazambuga et al., 1998; Gold, 2000; Kiggundu et al., 2003; Ocan et al., 2008). In particular, Fusarium wilt of banana (FWB) is one of the most devastating diseases affecting banana plants. The Tropical race 4 (TR4) of FWB caused by the soil-borne fungus *Fusarium oxysporum* f. sp. *cubense* (*Foc*) has put major constraints on global banana productions (Ordonez et al., 2015; van Westerhoven et al., 2022). The monoculture practice, the use of contaminated materials for planting and the lack of viable long-term solutions thereof, all contribute to the exacerbation of the diseases that are especially taxing on small holder farmers.

Advancing banana breeding by exploiting the genetic diversity of diploid breeding parents is key to improving polyploid hybrid bananas. For example, differential susceptibility to Fusarium wilt of banana (*Foc* TR4) has been described for key banana germplasm (Kung’u and Jeffries, 2001; Zuo et al., 2018; Arinaitwe et al., 2019; Ndayihanzamaso et al., 2020). Genetic variation in fruit quality traits, such as parthenocarpy or nutrient content, can only be accessed through trait introgression from diploid breeding material (Sardos et al., 2016a). Commercially, bananas are sold as triploids, but breeding triploid material is a complicated process that requires crossing fertile diploids to generate tetraploids, crossed again to diploids to produce secondary triploids (Brown et al. 2017; Batte et al. 2019). In addition, many large structural rearrangements have been characterized in banana breeding germplasm, which pose complications for introgressing key traits through recombination (Marin, et al., 2017; Němečková et al., 2018; Martin, et al., 2020a; Martin, et al., 2020b; Šimoníková et al., 2020). Genomics-guided breeding methods can help address the above challenges by enabling variant discovery, genetic mapping, and genomic selection to fast-track important traits (Uwimana et al., 2021; Chen et al., 2023a; Chen et al., 2023b; Bayo et al.; 2024). Accessing genomics-guided germplasm improvement requires investment in foundational genomics resources for banana.

To facilitate efforts to improve diploid banana breeding in quality and resistances against diseases and pests, an international team led by Bayer Crop Science, IITA, and the University of Queensland, through the TR4 Alliance, has made a genomics initiative to sequence the genomes of a collection of diploid *Musa* spp. and cultivars important for IITA’s breeding program in Africa. These include ‘Fu Des’ (ITC0939) and ‘Manameg Red’ (ITC0966) from *M. acuminata* spp. *banksii*; ‘Pahang’ (ITC0609), AA cv. ‘Rose’ (ITC0712), and ‘Pisang Lilin’ (ITC1121) from *M. acuminata* ssp. *malaccensis*; ‘Maia Oa’ (ITC0728) from *M. acuminata* ssp. *zebrina*; ‘Pisang Tongat’ (ITC0063) from *M. acuminata* ssp. *errans*. The selected germplasm span four key *M. acuminata* subspecies and are potential progenitors of the important banana cultivars ‘Matooke’, ‘Mchare’, Cavendish and other plantains (Němečková et al., 2018; Sardos et al., 2022; Li et al., 2024).

In this study, we used PacBio high-fidelity reads to generate haplotype resolved reference genomes for these phenotypically and taxonomically diverse *Musa* diploids. With these assemblies, we aim to increase the diversity and overall number of genome assemblies of key diploid banana parent lines. The genome data presented here complement the existing banana genome resources with complete genome sequences of agronomically-improved diploid hybrid parents. Given the undeterred spreading of the TR4 pandemic, threatening the major banana producing regions of the world (Martínez et al., 2023), this genomic resource will allow us to dissect TR4 resistance in divergent genotypes. This resource will enable fast tracking of resistance in conventional breeding through genomic prediction as well as gene editing of resistance traits as well as traits relating to fruit quality in breeding lines derived from these progenitors.

## Results

### Sequencing and assembly of seven high-quality banana genomes

High-molecular weight genomic DNA was extracted from cigar leaves of the seven banana cultivars in this study (**Table 1; Figure 1**) and sequenced on a PacBio Sequel IIe producing high fidelity (HiFi) reads (16.6-29.4 Gigabases (Gb) of sequence for each germplasm). The depth of coverage for the haplotype-resolved assemblies were in the range of 17 to 27x per assembly (**Table 2**). The 14 partially phased haplotype assemblies comprised of 247 to 1539 contigs, with the total assembly sizes ranging from 493 – 586 Megabases (Mb) (**Table 3**). The size of the DH-Pahang genome was estimated by flow cytometry at 523 Mb (D’Hont et al., 2012) and the latest long read assembly had a cumulative size of 485 Mb (Belser et al., 2021). Our haplotype phased ‘Pahang’ assemblies were comparably sized at 499 and 492 MB whereas the rest of the haplotype phased diploid assemblies had cumulative sizes between 506 Mb and 586 Mb, falling in the range of 94 to 112 % of the estimated genome size of ‘DH-Pahang’. All haplotype phased assemblies were comprised of 600 contigs or less with the number of N90 contigs ranging from 47 to 157. Only the assemblies for ‘Pisang Tongat’ and ‘Fu Des’ accessions had more contigs and were less contiguous than the rest of the collection. Furthermore, both ‘Pisang Tongat’ and ‘Fu Des’, as well as cv. ‘Rose’ and ‘Pisang Lilin’ had high heterozygosity rates at 1.47%, 1.32%, 1.80% and 1.50%, respectively (**Table 4**).

**Table 1.** The seven accessions sequenced in this study and their accession numbers, taxonomic classification, geographical location where these lines are likely originated from. “Edible” means the fruit is parthenocarpic; “wild” means the fruit only develop after pollination and will have seeds.

| Accession number | Accession name | *Musa acuminata* subspecies | Geographical range<sup>1</sup> | Edibility |
| --- | --- | --- | --- | --- |
| ITC0712 | AA cv. Rose | *malaccensis* | Malaysia<sup>2</sup> | Edible |
| ITC0609 | Pahang | *malaccensis* | Malaysia: Peninsular<sup>3</sup> | Wild |
| ITC1121 | Pisang Lilin | *malaccensis* | Malaysia, Indonesia<sup>4</sup> | Edible |
| ITC0939 | Fu Des | *banksii* | Papua New Guinea.<br>Australia: Queensland.<br>Samoa<sup>2</sup> | Edible |
| ITC0966 | Manameg Red | *banksii* | Papua New Guinea.<br>Australia: Queensland.<br>Samoa<sup>5</sup> | Edible |
| ITC0728 | Maia Oa | *zebrina* | Java/Indonesia<sup>6</sup> and<br>Hawaii<sup>7</sup> | Wild |
| ITC0063 | Pisang Tongat | *errans* | Philippines<sup>8</sup> | Some pulp |

<sup>1</sup> Based on a compilation of published taxa (Sachter-Smith, 2023).
<sup>2</sup> A derivative of *Musa acuminata* ssp. *malaccensis*. AA cv. 'Rose' and its somaclone 'Gu Nin Chiao', also known as IDN110, are frequent genitors in world breeding programmes due to the resistance and fruit quality traits they carry (Bakry et al., 2021).
<sup>3</sup> Taxa data provided by Simmonds N.W., 1957. Unique traits include 'Bracts red, non-imbricate' (Sachter-Smith, 2023).
<sup>4</sup> A derivative of *Musa acuminata* ssp. *malaccensis*. 'Pisang Lilin' from the Pisang Lilin subgroup of diploids with AA genomes (Ploetz et al., 2007).
<sup>5</sup> Taxa data provided by Mueller F.J.H. von, 1864. Unique traits include 'Pendent bunch, bend in rachis, large brown blotches on pseudostem' (Sachter-Smith, 2023).
<sup>6</sup> Originated from Indonesia: Java (Nasution, 1991) with unique traits such as 'leaves with red-brown blotches, fruits not hairy' (Sachter-Smith, 2023).
<sup>7</sup> In Hawai'i, this type of seeded variety was introduced as a medicinal plant and plants with these phenotypic descriptive is called 'Maia Oa' (Ploetz et al., 2007).
<sup>8</sup> Taxa data provided by Valmayor R.V., 2001. A unique trait includes 'Bracts green, large bunches up to 26 hands' (Sachter-Smith, 2023).

**Table 2.** Statistics of the HiFi libraries.

| Accession name | Total number of reads | Total bases | Average read length | N50 read length | Calculated depth of coverage of diploid genome<sup>1</sup> |
| --- | --- | --- | --- | --- | --- |
| AA cv. Rose | 1,913,537 | 29,271,776,776 | 15,297 | 16,504 | 26 |
| Pahang | 1,060,907 | 16,580,962,367 | 15,629 | 16,661 | 17 |
| Pisang Lilin | 1,700,965 | 22,758,223,451 | 13,379 | 14,182 | 21 |
| Fu Des | 2,285,116 | 19,738,030,518 | 8,608 | 8,583 | 17 |
| Manameg Red | 1,448,673 | 17,945,069,754 | 12,387 | 12,804 | 17 |
| Maia Oa | 1,983,197 | 29,413,041,262 | 14,831 | 15,876 | 27 |
| Pisang Tongat | 2,508,435 | 25,685,496,835 | 10,239 | 10,230 | 24 |

<sup>1</sup> Total bases of reads divided by sum of total bases of both haplotype phased assemblies.

**Table 3.** Genome assembly quality statistics. The assembly statistics for each haplotype assembly are indicated as hap1 | hap2 in each cell.

| Accession name | Number of contigs | Total bases | N50 | Number of N50 contigs | N90 | Number of N90 contigs |
| --- | --- | --- | --- | --- | --- | --- |
| AA cv. Rose | 430 \| 254 | 566Mb \| 544Mb | 10.46Mb \| 9.66Mb | 17 \| 20 | 2.41Mb \| 1.84Mb | 57 \| 64 |
| Pahang | 541 \| 247 | 499Mb \| 493Mb | 12.76Mb \| 11.66Mb | 13 \| 13 | 1.62Mb \| 2.05Mb | 50 \| 47 |
| Pisang Lilin | 625 \| 311 | 540Mb \| 506Mb | 7.36Mb \| 8.33Mb | 24 \| 19 | 1.44Mb \| 1.87Mb | 91 \| 70 |
| Fu Des | 1,539 \| 1,020 | 586Mb \| 544Mb | 1.60Mb \| 1.79Mb | 91 \| 71 | 0.28Mb \| 0.30Mb | 411 \| 357 |
| Manameg Red | 829 \| 485 | 539Mb \| 523Mb | 5.14Mb \| 3.74M | 31 \| 42 | 0.80Mb \| 0.82Mb | 136 \| 154 |
| Maia Oa | 603 \| 401 | 537Mb \| 531Mb | 5.01Mb \| 4.05Mb | 33 \| 36 | 0.69Mb \| 0.76Mb | 154 \| 157 |
| Pisang Tongat | 1,465 \| 569 | 554Mb \| 512Mb | 3.79Mb \| 4.08Mb | 41 \| 34 | 0.49Mb \| 0.77Mb | 184 \| 148 |

**Table 4.** The estimated and assembled genome sizes. Genome size estimates derived from k-mer analysis, as in Methods. The assembly statistics for each haplotype assembly are indicated as hap1 | hap2 in each cell where relevant. The heterozygosity rate was calculated using ‘GenomeScope2.0’ (Ranallo-Benavidez et al., 2020).

| Accession name | Estimated haploid genome (bp) | Assembly sizes (bp) | % difference to the estimated size | Heterozygosity (%) |
| --- | --- | --- | --- | --- |
| AA cv. Rose | 540,120,967 | 566,453,526 \| 544,298,584 | 4.88 \| 0.77 | 1.80 |
| Pahang | 523,216,062 | 499,014,147 \| 492,921,427 | -4.63 \| -5.79 | 0.49 |
| Pisang Lilin | 596,438,694 | 540,429,291 \| 506,266,108 | -9.39 \| -15.12 | 1.50 |
| Fu Des | 594,313,610 | 586,153,264 \| 544,383,587 | -1.37 \| -8.40 | 1.32 |
| Manameg Red | 628,597,152 | 538,694,546 \| 523,470,394 | -14.30 \| -16.72 | 0.45 |
| Maia Oa | 574,587,557 | 536,922,034 \| 531,126,841 | -6.56 \| -7.56 | 0.42 |
| Pisang Tongat | 594,128,672 | 554,270,387 \| 512,271,382 | -6.71 \| -13.78 | 1.47 |

**Figure 1.**
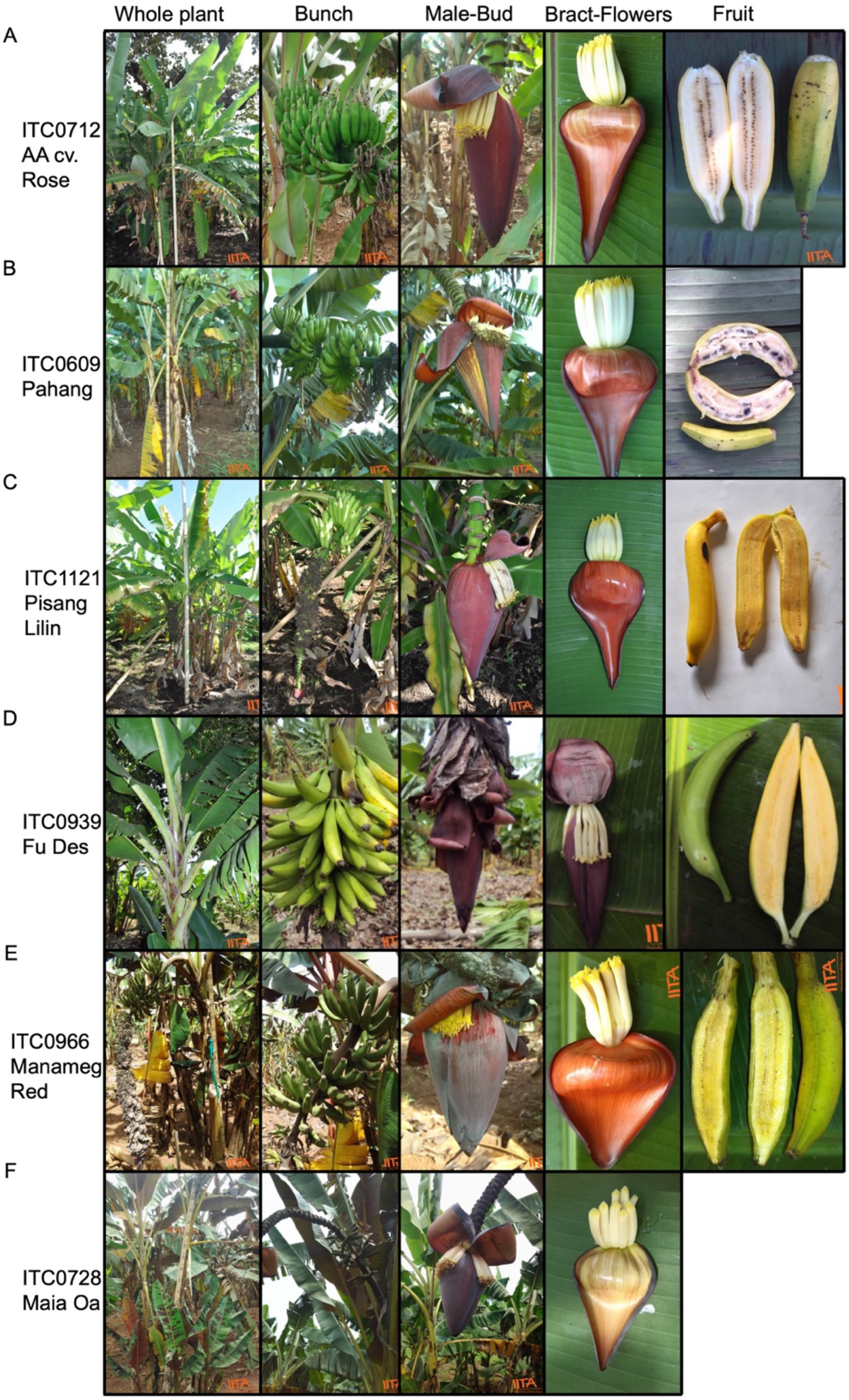
Phenotypic characteristics of six accessions sequenced in this study (Pisang Tongat not shown). Images downloaded from https://www.musabase.org in February 2024.

The HiFi reads were assembled with ‘hifiasm’ (Cheng et al., 2021), and produced two partially haplotype-resolved assemblies for each germplasm. Hi-C paired-end reads will be used in the future to correct haplotype switch and produce fully phased genomes for these accessions.

Assessing telomeric repeats at the ends of the assembled chromosomes showed that most chromosomes have telomeres placed on both the 5’ and 3’ ends, supporting the completeness of the assemblies (**Table 5**).

**Table 5.** Summary of telomeres detected in *de novo* partially phased genome assemblies for each cultivar. The telomeres detected for each haplotype assembly are indicated as hap1 | hap2 in each cell. In haplotype assemblies where both telomeres were not found, the result is indicated in bold text.

|  | AA cv. Rose | Pahang | Pisang Lilin | Fu Des | Manameg Red | Maia Oa | Pisang Tongat |
| --- | --- | --- | --- | --- | --- | --- | --- |
| *chr1* | **3' \| 3'** | **5' \| 5'** | both \| **3'** | both \| both | **5' \| both** | both \| both | both \| **3'** |
| *chr2* | both \| both | both \| both | both \| both | both \| both | both \| both | both \| both | both \| both |
| *chr3* | **5' \| none** | both \| both | both \| both | both \| both | **5' \| 5'** | **3' \| none** | both \| **5'** |
| *chr4* | both \| both | both \| both | **3'** \| both | both \| both | both \| both | both \| both | both \| both |
| *chr5* | **5' \| both** | both \| **5'** | both \| both | both \| both | **3' \| 3'** | both \| both | both \| both |
| *chr6* | **3' \| 3'** | both \| both | **5'** \| both | both \| **3'** | both \| **3'** | **5'** \| both | **3' \| 5'** |
| *chr7* | both \| both | both \| both | both \| both | **5'** \| both | both \| both | both \| both | both \| both |
| *chr8* | both \| both | both \| both | both \| **3'** | both \| both | both \| **none** | both \| both | both \| both |
| *chr9* | both \| both | both \| both | both \| both | both \| **5'** | both \| both | **5'** \| both | **3' \| 3'** |
| *chr10* | both \| **5'** | **5'** \| both | **5'** \| both | both \| **5'** | **5' \| 3'** | both \| both | **5'** \| both |
| *chr11* | both \| both | both \| both | **3'** \| both | **3' \| 3'** | **5' \| 3'** | both \| **none** | both \| both |

### K-mer analysis to estimate the genome size and heterozygosity rate

K-mer analysis was used to estimate the genome size using long read datasets (Ranallo-Benavidez et al., 2020). ‘Genomescope’ ver2.0 was used to estimate heterozygosity while a custom analysis on k-mer frequency and counts was used to estimate genome size of each assembly (Methods).

The estimated genome size of each assembly was in the range of 523 to 628 Mb, which was larger than the actual sizes of these assemblies (except for AA cv. ‘Rose’; **Table 4**). The relative heights of the peaks in a k-mer spectrum provided an estimate to the heterozygosity level of each accession (Ranallo-Benavidez et al., 2020) (Methods). The diploid assemblies had a heterozygosity range of 0.42-1.80%, which was in the range of expected heterozygosity for the sequenced material.

### Gene model annotations

Repeat-masking detected regions occupying 48.9% to 58.6% of the genome sequence that contain repetitive elements across all haplotype resolved assemblies. Out of these regions characterised, long terminal repeat elements occupied the majority at 39% to 51.3% of the genome sequence, whereas DNA elements and simple repeats respectively occupied 5.1% to 10.5% and 1% to 1.3% of the genome sequence. These results are comparable to the repetitive elements annotated totalling 52.6% of the genome in DH-Pahang (Belser et al., 2021).

‘GeMoMa’, a homology-based gene annotation program (Keilwagen et al., 2019), was used to annotate the haplotype resolved assemblies using gene models in five other *Musa* reference genomes. Firstly, reference genomes and annotations for *M. acuminata* Pahang v4, *M. beccarrii* v1, *M. textilis* (abaca) v1, *M. schizocarpa* v1, *M. troglodytarum* v1, *M. balbisiana* DH-PKW v1, *M. itinerans* v1, *M. acuminata* spp. *banksii* v2, *M. acuminata ssp. burmannica* v1 and *M. acuminata* ssp. *zebrina* v1 were retrieved from Banana Genome Hub (BGH) (https://banana-genome-hub.southgreen.fr/ accessed on 19 April 2024) and were subject to protein BUSCO analysis (Manni et al., 2021). Out of these references, five reference genomes showed a protein based BUSCO higher than 90%, including *M. acuminata* Pahang v4 (97.1%), *M. beccarrii* v1 (94.8%), *M. textilis* (abaca) v1 (92.7%), *M. schizocarpa* v1 (91.6%), *M. troglodytarum* v1 (90%). These genomes were then used as reference species for ‘GoMoMa’ since its prediction quality depends on the quality of the target genome sequence (Keilwagen et al., 2019).

Transcript sets from these reference genomes were retrieved from BGH (https://banana-genome-hub.southgreen.fr/ accessed on 19 April 2024) and were used to predict gene models for each of the 14 haplotype resolved assemblies using ‘GeMoMa’. This resulted in the annotation of approximately 38,000 gene models in each new genome (**Table 6**). The number of gene models was higher than 36,979 genes annotated in DH-Pahang v4 (Belser et al., 2021). Despite not having RNA-seq reads from each accession to provide evidence in building transcript models, the BUSCO scores for the predicted amino acid sequences generated by ‘GeMoMa’ annotations showed 98.0-98.9% genome completeness for each of the 14 haplotype resolved genomes, in comparison to a 96.0% genome completeness assessed for the DH-Pahang v4 assembly.

**Table 6.** Counts of gene model and transcript annotations and BUSCO Scores for each assembly obtained from ‘GeMoMa’ (see Methods).

| Accession name | Haplotype | Annotated feature count<br>(scaffolded contigs only) |  |  | BUSCO calculated based on protein sequence |  |  |  |  |
| --- | --- | --- | --- | --- | --- | --- | --- | --- | --- |
|  |  | gene | mRNA | putative NLRs | Complete | Single copy | Multi-copy | Fragmented | Missing |
| AA cv. Rose | hap1 | 38625 | 45180 | 142 | 98.1 | 78.8 | 19.3 | 0.7 | 1.2 |
| AA cv. Rose | hap2 | 39046 | 45605 | 113 | 98.3 | 77.8 | 20.5 | 0.7 | 1.0 |
| Pahang | hap1 | 39673 | 46302 | 144 | 98.8 | 80.7 | 18.1 | 0.6 | 0.6 |
| Pahang | hap2 | 39683 | 46326 | 129 | 98.8 | 80.7 | 18.1 | 0.6 | 0.6 |
| Pisang Lilin | hap1 | 38987 | 45610 | 136 | 98.7 | 79.7 | 19.0 | 0.7 | 0.6 |
| Pisang Lilin | hap2 | 38884 | 45514 | 137 | 98.7 | 80.4 | 18.3 | 0.6 | 0.7 |
| Fu Des | hap1 | 38307 | 44842 | 138 | 98.2 | 78.8 | 19.4 | 0.8 | 1.0 |
| Fu Des | hap2 | 38292 | 44872 | 142 | 98.2 | 79.3 | 18.9 | 0.8 | 1.0 |
| Manameg Red | hap1 | 38037 | 44625 | 114 | 98.5 | 79.8 | 18.7 | 0.8 | 0.7 |
| Manameg Red | hap2 | 37929 | 44457 | 105 | 98.5 | 80.3 | 18.2 | 0.8 | 0.7 |
| Maia Oa | hap1 | 37991 | 44518 | 162 | 98.5 | 80.3 | 18.2 | 0.9 | 0.6 |
| Maia Oa | hap2 | 38018 | 44575 | 165 | 98.3 | 80.2 | 18.1 | 1.0 | 0.7 |
| Pisang Tongat | hap1 | 38470 | 45077 | 112 | 98.6 | 80.3 | 18.3 | 0.7 | 0.7 |
| Pisang Tongat | hap2 | 38104 | 44655 | 161 | 98.0 | 79.7 | 18.3 | 0.8 | 1.2 |

Finally, we assessed the abundance of disease resistance genes particularly those encoding nucleotide-binding and leucine-rich repeat (NLR) proteins in our haplotype resolved assemblies. These immune receptors play important roles in plant surveillance of invading pathogens during infection and intracellularly triggers an immune response (Jones and Dangl, 2006). An NLR prediction software, namely the ‘NLR-annotator’ was used to annotate loci associated with protein motifs usually associated with NLRs (Steuernagel et al., 2020). We identified 105 to 165 loci containing NLR motifs in the haplotype resolved assemblies (**Table 6**). Variations in the number of NLR loci were evident in pairs of some of the haplotype resolved genomes, including ‘Pisang Tongat’, which had 112 NLR loci in one and 161 in the other haplotype resolved genome (**Table 6**).

### Gene synteny analysis and detection of rearrangements

Gene order and orthology amongst all haplotype resolved genomes were examined at the chromosome level using the software ‘GENESPACE’ (Lovell et al., 2022). The syntenic relationships were visualized based on the physical position of all genes across all haplotype phased genomes. Overall, conservation of gene content and order was evident amongst these assemblies at the chromosome level, and some local rearrangements and inversions were also detected (Figure 2).

**Figure 2.**
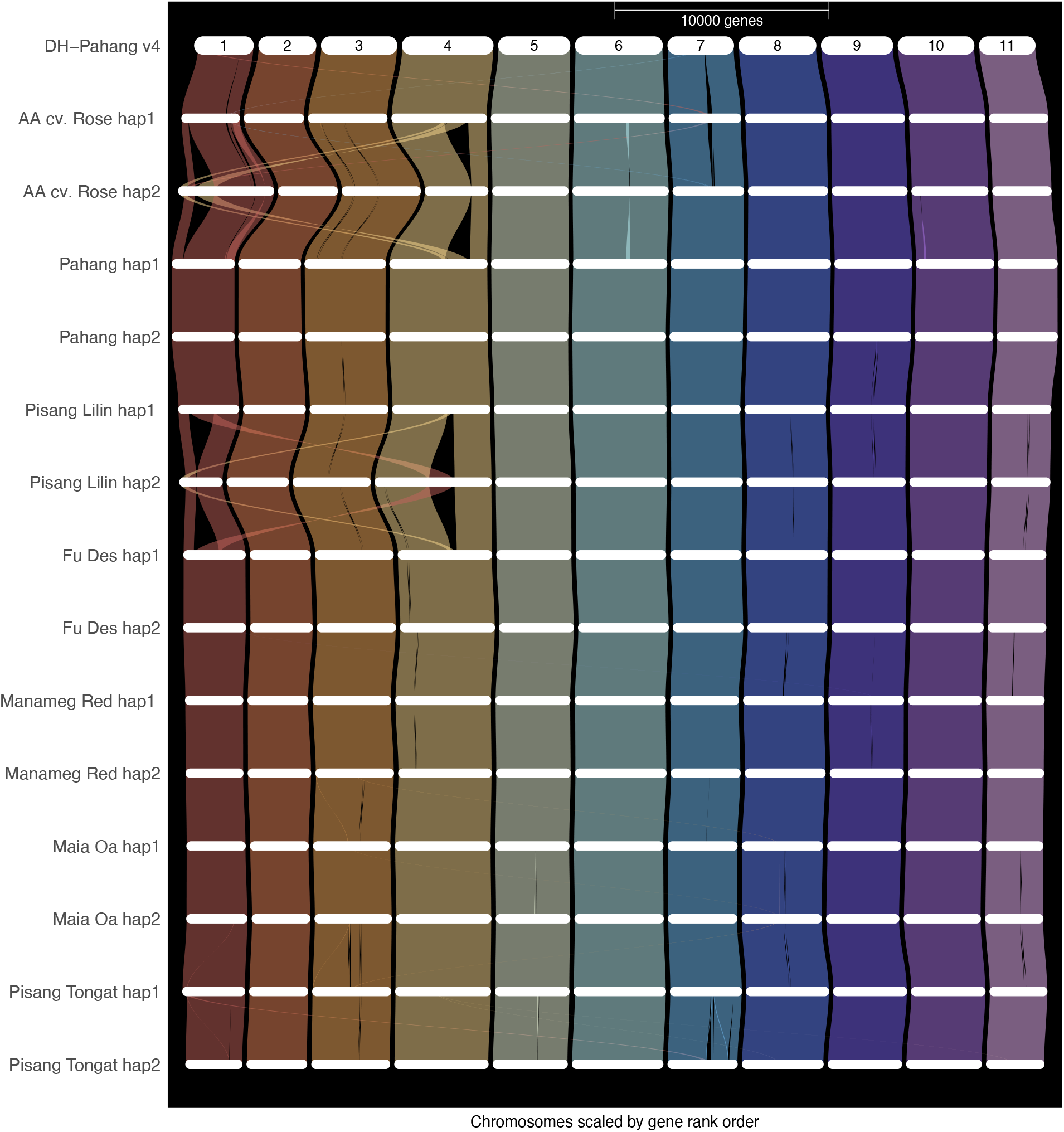
GENESPACE plot of the 14 partially phased assemblies reported here and DH-Pahang v4.

There was no structural discordance indictive of incorrectly positioned contigs at the chromosome level. Further sequencing to detect long-range interactions (particularly Hi-C or ultra long reads) will support scaffolding potentially address some of the finer discrepancies, particularly with respect to some of the well-known translocation events (Baurens et al., 2018; Dupouy et al., 2019; Martin et al., 2020a). A specific reciprocal translocation involving chromosome 1 and chromosome 4 was previously reported and characterised in ‘Pisang Lilin’ (Martin et al., 2017; Šimoníková et al., 2020) and is detected in the initial haplotype assemblies for ‘Pisang Lilin’ presented here.

## Discussion

In this study, we produced preliminary genome drafts of seven diploid (AA) banana accessions. Unlike the B genome progenitor *Musa balbisiana* which has its genome size estimated at 534-540 Mb using the 2C nuclear DNA content (Lysak et al., 1999), the AA genome appeared to be relatively larger compared to *M. balbisiana*, with wild relatives and cultivated diploids estimated at 591-615 Mb (Lysak et al., 1999), 580-640 Mb (Kamaté et al., 2001), and 523 Mb (D’Hont et al., 2012). ‘Pahang’ is the parent to ‘DH-Pahang’ or ‘CIRAD930’ and these have been shown to be genetically similar accessions in genotyping studies (Christelová et al., 2017; Ruas et al., 2017). Our haplotype resolved ‘Pahang’ assemblies are 492 and 499 Mb in size, and has a predicted size of 523 Mb, both of which are consistent with the characterisation of ‘DH-Pahang’ in previous studies. The other two *M. acuminata* ssp. *malaccensis* derived diploids, AA cv. ‘Rose’ and ‘Pisang Lilin’, had haplotype resolved assembly sizes of 544 and 566 Mb, and 506 and 540 Mb, respectively, which are larger than ‘DH-Pahang’ but smaller than the size of other AA diploids of *M. acuminata* determined by flow cytometry in previous studies (Lysak et al., 1999; Kamaté et al., 2001).

The two *M. acuminata* ssp. *banksii* derivatives, ‘Fu Des’ and ‘Manameg Red’ had haplotype resolved genome sizes of 544 and 586 Mb, and 523 and 538 Mb, respectively, which are considerably smaller than the estimated genome sizes of three distinct types of the wild species of *M. acuminata* ssp. *banksii* at 580-630 Mb (Kamaté et al., 2001). *M. acuminata* ssp. *zebrina* ‘Maia Oa’ produced 531 and 536 Mb haplotype resolved assemblies, which are shorter than the Illumina short read-based assembly of the same line at 623 Mb (Rouard et al., 2018) but comparable to an Oxford Nanopore Technology (ONT) long read assembly of *M. acuminata* ssp. *zebrina* v2.0, totalling 558 Mb in cumulative size (Li et al., 2024). Finally, *M. acuminata* ssp. *errans* derived ‘Pisang Tongat’ had haplotype resolved genome sizes of 512 and 554 Mb, which is smaller than the estimated size of *M. acuminata* ssp. *errans* at 606 Mb (Doležel et al., 1994). Additional sequencing data is expected to improve the phasing of these assemblies, which may alter the overall size of the haplotype resolved assemblies.

The cost of DNA sequencing has declined in the era of third generation long-read sequencing technologies, facilitating genome assemblies for large plant polyploid genomes (Sahu and Liu, 2023). However, technical hurdles still exist in producing fully phased plant genomes (Delorean et al., 2023). When an accession exhibits a high level of heterozygosity such as the ones presented in this study, the chance that the resulting assemblies contain chimeric regions of the individual’s two haplotypes is high. These chimeric regions may provide misleading information in downstream analyses in gene discovery. Bionano optical maps, Chromatin conformation capture or Hi-C sequencing, ultra-long DNA sequencing reads, and genetic maps have been used as physical or genetic information to determine which assembly contigs or haplotigs group together to form a chromosome (Sedlazeck et al., 2018; Belser et al., 2021; Cochetel et al., 2023; Liu et al., 2023). Hi-C has been widely used to aid chromosome scale scaffolding in haplotype resolved plant genome assemblies (Šimková et al., 2024). In *Musa* spp., the latest genomes that have been fully haplotype-resolved have seen an increasing use of Hi-C as the scaffolding tool to resolve haplotypes in diploid and triploid cultivars (Huang et al., 2023; Liu et al., 2023; Li et al., 2024). As evident in our assemblies, some of the translocation events will need to be distinguished from assembly errors by of performing Hi-C or other long-range sequencing validation in our assemblies.

One of the haplotype phased assemblies in ‘Pisang Lilin’ and AA cv. ‘Rose’ appeared to carry reciprocal translocations involving chromosome 1 and 4. These translocations were previously reported in ‘Pisang Lilin’ (Martin et al., 2017; Šimoníková et al., 2020). The same translocations were also confirmed in ‘Cavendish’, ‘Gros Michel’ and ‘Mchare’ cultivars (Šimoníková et al., 2020). The precise location and the structure of the translocated fragments requires additional analysis to resolve in our assemblies.

In banana breeding, efforts have been made in the last decades to introgress traits relating to disease resistance in Africa (Lorenzen et al., 2010). This has been hindered by the long generation times of growth, fertility issues with the polyploids, and limited genetic diversity captured by the core *Musa* collection in breeding (Ortiz and Swennen, 2014). Often, banana breeding pipelines relied on diploids to increase genetic diversity of East African highland bananas through recurrent selection of important traits. Thus, improving the quality of diploids is an important activity for breeding. Here, we selected a set of diploids important in the IITA breeding program for sequencing. These diploids appear to show genetic diversity in whole genome diversity analyses (Sardos et al., 2016b; Christelová et al., 2017).

These assemblies when phased and structurally validated, will support banana breeding and trait discovery research programs. They will also serve as a key step in broadening genome sequence information for this important crop. Next steps for this project include generating more long-range interaction data to fully phase these assemblies and support the translocation events detected so far. Fully phased assemblies will also enable a closer examination of the TR4 resistance loci (Ahmad et al., 2020; Chen et al., 2023a; Chen et al., 2023b) and other key trait genes (Li et al., 2024).

## Materials and Methods

### Source Material

The seven *Musa* accessions sequenced in this study were as listed in Table 1. Lyophilized young leaves were shipped to Bayer Crop Science Precision Genomics Unit (St. Louis, MO, USA) for DNA extraction, library preparation and Pacific Biosciences HiFi sequencing.

### Extraction, Library Preparation & Sequencing

All lyophilized tissue was ground in liquid nitrogen to fine powder. To approximately 1 gram of leaf powder, 20 mL of pre-heated (65°C) modified Cetyltrimethylammonium bromide (CTAB) extraction buffer [2% CTAB (H-5882, Merck, Darmstadt, Gerrmany); 100 mM Tris-HCl (pH8.0); 20 mM Ethylenediaminetetraacetic acid (EDTA, pH8.0); 1.4 M Sodium Chloride, freshly added 1% Polyvinylpyrrolidone and 2% (v/v) Tween-20] was added and vortexed gently until complete resuspension. The resuspended samples were incubated at 65ºC for 45 minutes with gentle mixing every 10 minutes. After incubation, 20 μl of Proteinase K (P8107S, New England Biolabs, Ipswich, MA, USA) was added and incubated at 60°C for 15 minutes with gentle mixing every 5 minutes. DNA was purified using equal volumes of Phenol Chloroform Isoamyl (24:24:1) followed by Chloroform:Isoamyl alcohol extraction (24:1). DNA was purified by precipitation of aqueous phase by equal volume isopropanol and centrifugation.

Integrity and concentration of isolated genomic DNA was measured by Qubit (Thermo Fisher, Waltham, MA, USA), Nanodrop (Thermo Fisher, Waltham, MA, USA) and Femto Pulse system (Agilent, Santa Clara, CA, USA). 10 μg of resuspended DNA was subjected to a bead cleanup step (Genfind v3.0, Beckman Coulter, Brea, CA, USA) and 5 μg of DNA were used for HiFi library preparation. All HiFi libraries were prepared using Pacific Biosciences SMRTbell® Express Template Prep Kit 3.0 and sequenced on the Pacific Biosciences Sequel IIe system to generate Circular Consensus (CCS) reads.

### K-mer analysis

For each of the seven banana accessions, a k-mer analysis was performed on the HiFi reads with ‘Jellyfish’ v.2.2.4 (https://github.com/gmarcais/Jellyfish; (Marçais and Kingsford, 2011)) using a k-mer size of 55 and a histogram of the k-mer frequency was generated. The computation was run on an AWS EC2 r5.12xlarge node (48 vCPUs and 380Gb RAM). The output of jellyfish ‘histo’ program is a data set with counts of k-mers at given frequencies; this table was then used to estimate genome sizes for each accession. The genome size for a given assembly was estimated as the product of the total number of k-mers times the k-mer size (55 in this case), divided by the frequency mode from k-mer distribution. The same k-mer frequency table was used as the input for ‘Genomescope’ heterozygosity rate estimation.

### Assembly with HiFi reads and pseudomolecule scaffolding

All seven banana genomes were sequenced on a Pacific Biosciences Sequel IIe system. The sequencing adaptors and multiplexing barcodes were removed with PacBio SMRTLink version 11.1 after the demultiplexing was finished.

Each of these seven genomes was assembled using ‘hifiasm’ v. 0.18.2-r467 (Cheng et al., 2022). The assembly process of each genome was run on an AWS EC2 r5.12xlarge node (48 vCPUs and 380Gb RAM). The version 0.18.2 of ‘hifiasm’ produces two sets of partially phased haploid contigs by default. The heterozygosity rates estimated by Genomescope2.0 (Ranallo-Benavidez, 2020) were predicted to be lower than 0.5% in ‘Pahang’, ‘Manameg Red’ and ‘Maia Oa’, therefore the default parameter of 0.55 for setting -s for ‘hifiasm’ was used. The estimated heterozygosity rates of AA cv. ‘Rose’, ‘Pisang Lilin’, ‘Fu Des’, and ‘Pisang Tongat’ were all larger than 1.3%; therefore, the option -s for ‘hifiasm; was set to 0.5, instead of 0.55, to avoid unbalanced sizes of partially phased haploid genomes.

To scaffold contigs into pseudomolecules, a reference-based scaffolding method, ‘RagTag’ (v2.1.0) (Alonge *et al*., 2022), was adopted. The ‘scaffold’ tool was used align the ‘hifiasm’ contigs with the DH-Pahang v4 reference (Belser *et al*., 2021), and subsequently clustered contigs together into groups using the corresponding physical information from the 11 *Musa acuminata* chromosomes and mitochondrionrial genome of DH-Pahang v4. Then the contigs of each chromosome group were ordered based on their alignment positions and joined to form continuous pseudomolecules. Unscaffolded contigs for each haplotype assembly were concatenated onto chr10000001 with 100 “N” bases inserted between subsequent contigs.

### Gene model annotation

Non-redundant transposable element libraries were created for each genome using ‘EDTA’(v2.0.0, --sensitive 0) (Ou et al., 2019). These libraries were then passed to ‘RepeatMasker’ (version open-4.0.9) (Smit et al., 2013) for repeat masking. ‘GeMoMa’ (v1.9) (Keilwagen et al., 2018) was used to map gene models from give public genome datasets: Musa_acuminata_pahang_v4, Musa_beccarii_1.0, Musa_textilis_abaca_1.0, Musa_schizocarpa_1.0, and Musa_troglodytarum (GFF3 and fasta files were downloaded from the Banana Genome Hub (https://banana-genome-hub.southgreen.fr/, accessed in March 2024). ‘GeMoMa’ annotation filter parameters were chosen to limit less conserved gene predictions (NaN(score) or score/aa>=4). BUSCO scores were calculated using the BUSCO tool suite version 5.5.0 with the database ‘liliopsida_odb10’ (creation date 2017-12-01) (Manni et al., 2021). The ‘NLR-Annotator’ (v2.1, default settings) was used to identify putative NLR loci (Steuernagel et al., 2020). ‘GENESPACE’ (v1.3.1, default settings) was used for gene synteny analysis (Lovell et al., 2022).

